# MicroPIPE: *An end-to-end solution for high-quality complete bacterial genome construction*

**DOI:** 10.1101/2021.02.02.429319

**Authors:** Valentine Murigneux, Leah W. Roberts, Brian M. Forde, Minh-Duy Phan, Nguyen Thi Khanh Nhu, Adam D. Irwin, Patrick N. A. Harris, David L. Paterson, Mark A. Schembri, David M. Whiley, Scott A. Beatson

## Abstract

Oxford Nanopore Technology (ONT) long-read sequencing has become a popular platform for microbial researchers; however, easy and automated construction of high-quality bacterial genomes remains challenging. Here we present MicroPIPE: a reproducible end-to-end bacterial genome assembly pipeline for ONT and Illumina sequencing. To construct MicroPIPE, we evaluated the performance of several tools for genome reconstruction and assessed overall genome accuracy using ONT both natively and with Illumina. Further validation of MicroPIPE was carried out using 11 sequence type (ST)131 *Escherichia coli* and eight publicly available Gram-negative and Gram-positive bacterial isolates. MicroPIPE uses Singularity containers and the workflow manager Nextflow and is available at https://github.com/BeatsonLab-MicrobialGenomics/micropipe.

## Background

Bacterial genome construction using short-read sequencing has historically been difficult, largely due to the abundance of repeat sequences which collapse during *de novo* assembly, resulting in breaks in contiguous sequence [1]. However, long-read sequencing technologies, such as Oxford Nanopore Technology (ONT) and Pacific Biosciences (PacBio), are able to traverse these repeats enabling complete bacterial genomes [2]. Long reads also present the opportunity to correctly place single nucleotide variants (SNVs), particularly across complex regions of the genome that require more genomic context than short reads can provide. The accessibility and affordability of the ONT MinION sequencing device has resulted in its widespread use globally, allowing researchers the autonomy to perform their own experiments much more rapidly than through external sequencing facilities [3]. However, bacterial genome construction continues to be problematic, especially for non-specialised researchers.

Numerous tools designed to address aspects of complete bacterial genome construction have been developed by both ONT and community users, however few pipelines exist that offer end-to-end construction of bacterial genomes. Currently, these include Katuali (ONT), CCBGpipe [4], ASA^3^P [5] and Bactopia [6]. Katuali is an ONT developed assembly pipeline implemented in Snakemake. It offers the user flexibility in software choice, but with limited guidance or rationale. While ASA^3^P and Bactopia are able to generate assemblies using nanopore data, overall these pipelines were not designed solely for *de novo* assembly and are more focused on reproducible and comprehensive downstream analysis. CCBGpipe is distributed via Docker and implements a series of python scripts to run Canu with Racon and Nanopolish. However, this pipeline performs Nanopore-only assembly (without Illumina) and was designed using Canu version 1.6, which is now several releases behind the current version (v2.1.1).

Substitution errors in nanopore reads have improved dramatically over recent years, from read accuracies of 60% [7] to the currently reported 95% for 1D reads using R9.4.1 flow cells [8]. While this is approaching that of Illumina (99.9%) [9] and PacBio (99%) [10], single nucleotide insertion/deletion (indel) errors remain problematic [11, 12]. Improvements in base-calling software (e.g. that account for methylation) and the introduction of the R10 pore have reduced these artefacts, but polishing nanopore assemblies with Illumina data has been generally required to achieve the highest quality possible [13].

With the rapid pace of ONT progression, development of new software and pipelines, or reappraisal of existing ones, has become an ongoing necessity. This has prompted the need for appropriate validation sets, to assess (or reassess) the accuracy of results. While simulated datasets provide an initial assessment of a tool’s ability, data generated from biological sources provide additional confidence in its real-world application, as has been developed previously using metagenomic communities [14, 15]. *Escherichia coli* sequence type (ST)131 represents a globally disseminated lineage that has been intensively studied as a result of its recent emergence, antibiotic resistance and link to human disease [16–18]. Extensive knowledge of both *E. coli* (as a species) and the ST131 lineage makes it an ideal dataset to use for software and pipeline validation. Additionally, the *E. coli* ST131 strain EC958 represents an extensively curated and highly accurate reference genome, having been sequenced on multiple occasions using PacBio, Illumina and 454 pyrosequencing [19].

Here we present our complete pipeline, MicroPIPE, for automated construction of high-quality bacterial genomes using software chosen by systematic comparison of the most popular tools currently available in the community. Validation of each pipeline stage was completed using the high-quality *E. coli* ST131 reference genome, EC958. Subsequent validation of the complete pipeline was performed using 11 previously characterised ST131 *E. coli* strains, for which completely assembled genomes were already available. Finally, we tested MicroPIPE on eight other publicly available bacterial isolates that had both a complete genome and associated raw nanopore sequencing data available. In all cases, we show that high-quality bacterial reference genomes can be achieved using MicroPIPE.

## Results

### Section 1: pipeline results and comparison to EC958 complete genome

The main goal of this study was to create a robust and easily applicable pipeline for the construction of high-quality bacterial genomes with minimal manual manipulations. To achieve this, we first evaluated the performance of commonly used software at each stage of bacterial genome construction using the high-quality EC958 genome (Accession: HG941718) as our standard for final genome accuracy. **Figure 1** shows a diagram of the whole workflow, indicating the software chosen for comparison at each stage. Nanopore reads for EC958 were generated on a multiplexed run of 12 using the rapid barcoding kit on an R9.4.1 flow cell.

**Figure 1:**
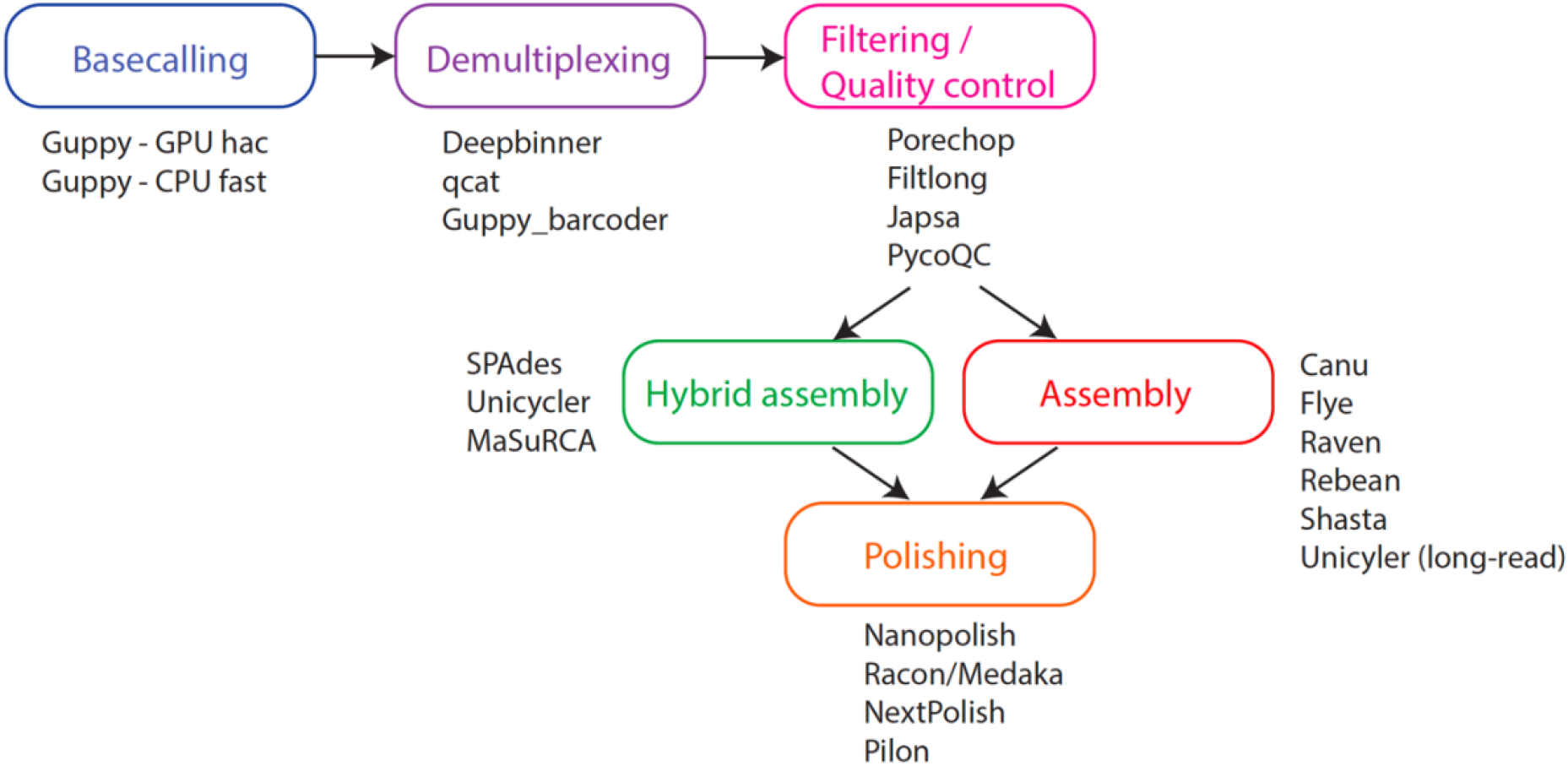
overall diagram of assembly stages and tool comparisons.

#### Basecalling

To evaluate basecalling, we tested Guppy using both the “fast” and “high-accuracy” modes, as well as the CPU vs. GPU configurations. When using Guppy v3.4.3 with the “high-accuracy” setting on GPU servers we generated reads with approximately 91.0% accuracy in 828.5 minutes (13.81 hours). Using the “fast” mode on CPUs, we were able to generate 88.9% accuracy in 2948.4 minutes (49.14 hours) **(Table 1)**. Testing the “high-accuracy” mode on a CPU server was unfeasible due to the time required for processing (fewer than 10% of reads completed basecalling in one week). Despite the lower per-read accuracy when using CPUs and the “fast” basecalling setting, the consensus quality of the overall finished genome (after assembly and polishing through MicroPIPE v0.8) was of comparable quality to that generated with the GPU and high-accuracy setting **(Table 1)**.

**Table 1:** Basecalling comparison: run-times, read accuracy and overall assembly accuracy.

| **Basecalling comparison:** | Guppy3.4.3_hac | Guppy3.4.3_fast | Guppy3.4.3_hac_<br>modbases | Guppy3.6.1_hac | Guppy3.6.1_hac_<br>modbases |
| --- | --- | --- | --- | --- | --- |
| *Run time (ms)* | 49,707,952 | 176,906,144 | 57,479,661 | 57,977,178 | 46,296,565 |
| *Run time (h)* | 13.81 | 49.14 | 15.96 | 16.10 | 12.86 |
| *GPU/CPU* | GPU | CPU | GPU | GPU | GPU |
| *Num callers* | 4 | 16 | 8 | 8 | 8 |
| *Average read percent identity* | 91.0 | 88.9 | 90.6 | 93.7 | 91.0 |
| *Mean read quality* | 11.4 | 10.4 | 11.3 | 13.3 | 11.4 |
| *Number of binned reads (qcat)* | 240,766 | 233,802 | 238,847 | 244,830 | 240,156 |
| **Final assembly comparison:** |  |  |  |  |  |
| *Assembly nucleotide identity (%)* | 99.99 | 99.99 | 99.99 | 99.99 | 99.99 |
| *Number of SNP (DNAdiff)* | 23 | 35 | 3 | 4 | 5 |
| *Number of indels (DNAdiff)* | 45 | 39 | 31 | 25 | 27 |
| *Assembly quality score (Pomoxis)* | 48.10 | 48.08 | 50.99 | 52.27 | 51.83 |
| *Mismatches per 100 kb (QUAST)* | 0.44 | 0.67 | 0.06 | 0.08 | 0.10 |
| *Indels per 100 kb (QUAST)* | 0.88 | 0.76 | 0.63 | 0.50 | 0.53 |

We also tested the effects of methylation and found that using the “high-accuracy” model with methylation-aware basecalling achieved a similar per-read accuracy (90.6%) to the “high-accuracy” only model. The final assembly, however, had fewer SNPs (3 vs. 23 originally) and indels (31 vs. 45 originally) compared to the reference standard **(Table 1)**.

#### Demultiplexing

For demultiplexing we tested three tools: Deepbinner, Guppy_barcoder and qcat. While Guppy and qcat rely on basecalled reads, Deepbinner uses the raw fast5 reads. As such, we compared the total numbers of binned reads after both basecalling and binning for each tool. Overall qcat was the fastest demultiplexer, and was able to bin 89% of reads, compared to 84% for Guppy_barcoder and 75% for Deepbinner **(Supplementary Figure 1)**. We prioritised read retention to maximise coverage of each genome. As such, qcat was chosen as the default demultiplexer for MicroPIPE. Following the recent depreciation of qcat, we have also provided Guppy_barcoder as an optional demultiplexer.

#### Filtering

Here we trialled two filtering tools: Filtlong and Japsa. Filtlong has the advantage of being versatile enough to filter based on a number of requirements, such as read length, quality, percentage of reads to keep and the option of using an external reference. Japsa primarily filters based on read length and quality. Read metrics after filtering using each tool are given in **Supplementary figure 2.** Overall, we found that filtering with Japsa retained more reads, but with a reduced N50 read length and median read quality compared to Filtlong. Both tools took an equivalent amount of time to run. For all downstream analysis we filtered reads using Japsa with a minimum average quality cut-off of Q10 and 1 kb minimum read length, although Filtlong would have been equally suitable. Both filtering tools are available as optional steps in micropipe.

#### Assembly

A number of tools have been designed for *de novo* assembly from long reads. Here we compared six popular assembly tools and evaluated speed, completeness (of the chromosome and plasmids, including circularisation) and correctness (i.e. nucleotide identity) based on the complete EC958 reference genome standard, which contains 1 chromosome (5,109,767 bp) and 2 plasmids (135,602 bp and 4080 bp). Parameters used for all assemblers are given in **Supplementary Dataset 1**.

Overall, we found that all assemblers constructed the chromosome and larger (~135 kb) plasmid **(Figure 2, Supplementary Table 2)**. Raven, Redbean and Shasta did not assemble the smaller ~4 kb plasmid. While Canu was able to assemble both plasmids, closer inspection found them to be much larger than expected (1.4x and 2x larger for the large and small plasmid, respectively) due to overlapping ends that required additional trimming. Interestingly, both Flye and Canu assembled a third, previously unidentified, small plasmid of ~1.8 kb in size. This small plasmid was only identified when the Flye “—plasmids” mode was selected (to rescue short unassembled plasmids) and when certain or no filtering parameters were applied to the reads prior to assembly (**Supplementary Table 3**). Comparison of this small plasmid to the Illumina data for the EC958 reference genome standard confirmed its presence and was likely missed in the original assembly.

**Figure 2:**
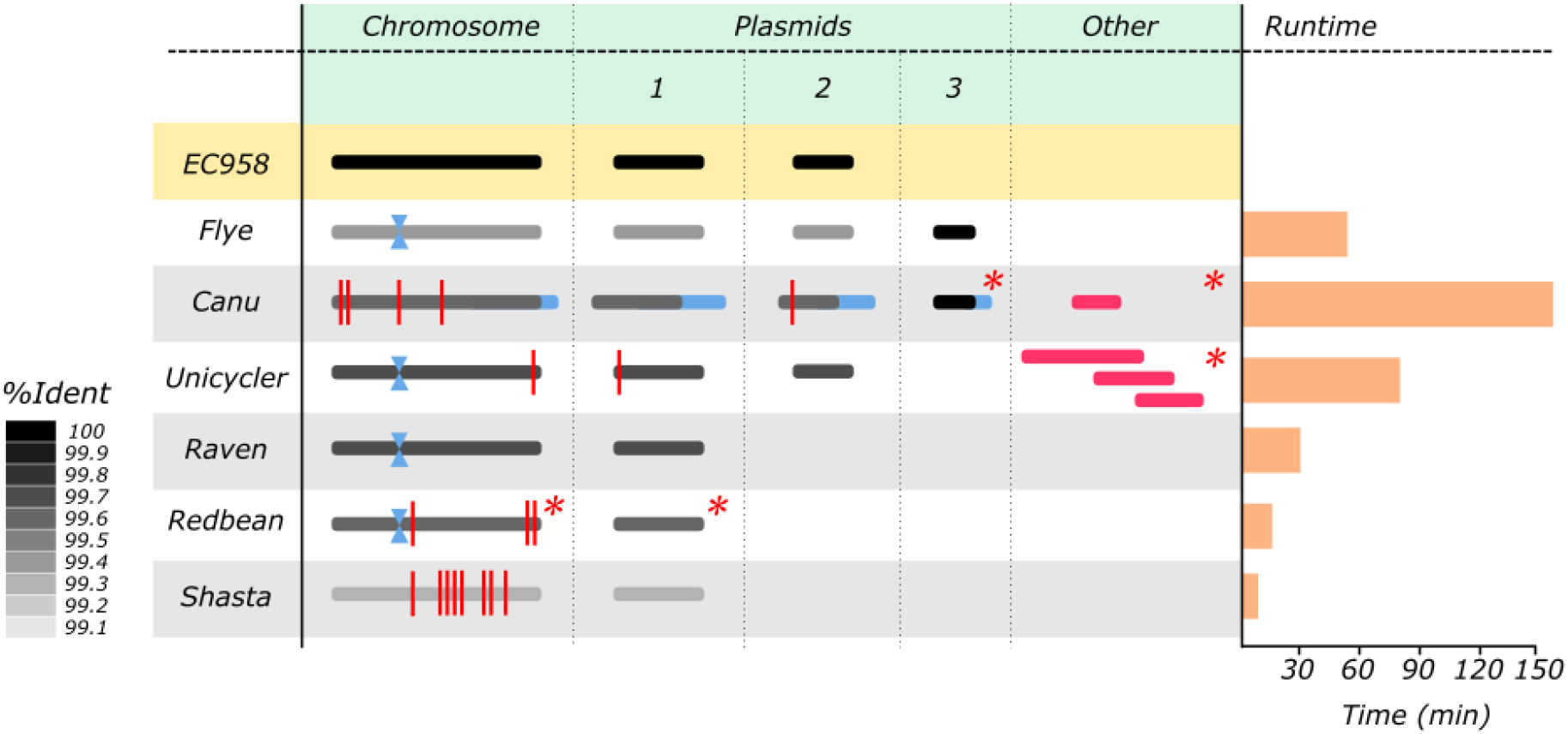
Assembly comparison: long horizonal bars represent contiguous sequences generated by each assembler. The chromosome and plasmids 1 and 2 are coloured according to their overall nucleotide identity when compared to the EC958 reference genome standard. The additional blue bars in the Canu plasmids represent the increased size of the plasmids from this assembler. Contigs that were not reported as circularised are marked with a red asterisk (*). Misassemblies are marked with a red vertical line at their approximate position. The phage tail protein inversion is marked with a blue hourglass.

For most *de novo* assemblies, a number of small (<4.5 kb) misassemblies were detected, mainly on the chromosome (**Figure 2**). This included a small inversion, which on closer inspection was found to be an invertible phage tail protein that has been characterised previously [19]. This inversion was found in the Flye, Unicycler, Raven and Redbean assemblies and was not counted as a misassembly due to its biological relevance.

Additional contigs were found in both Canu and Unicycler (long-read only mode). The three additional contigs produced by Unicycler all matched other parts of the EC958 reference genome standard (two on the chromosome, one on the larger plasmid). The additional contig in Canu matched part of the additional ~1.8 kb plasmid.

In terms of speed, Shasta, Redbean and Raven were the fastest assemblers, completing in less than 30 minutes. Of the remainder, Flye was four times faster than Canu and two times faster than Unicycler. The majority of contigs from all assemblers were reported as circularised upon assembly completion, with the exception of the additional contigs in Canu and Unicycler. Redbean did not generate circularisation information, although the chromosome and plasmid contigs could be circularised manually or using 3^rd^ party software following assembly. Overall, we found that Flye generated the best *de novo* assembly from long read data without the need for manual intervention.

#### Polishing

Polishing of assemblies generated using long reads is currently regarded as a necessity for ONT data due to high per-read errors that can persist through to the *de novo* assemblies [13]. Here we tested the polishing capabilities of three different tools (Racon/Medaka, NextPolish and Nanopolish) using nanopore long reads against the *de novo* assembly generated using Flye. We additionally tested polishing with Illumina short reads (NextPolish and Pilon), which have a higher basecall accuracy. Polishing was tested both independently (i.e., long read and short read separately) as well as sequentially (long read followed by short read polishing) to determine the best polishing protocol.

Overall, we found that polishing with Racon and Medaka (using long reads) followed by NextPolish (using short reads) achieved the most accurate assemblies **(Figure 3, Supplementary Table 4)**. Polishing using only long or short reads did not produce comparable levels of accuracy, therefore we emphasize the requirement of short read sequencing in parallel with Nanopore for high-quality complete genome assembly (as is already commonly done).

**Fig 3:**
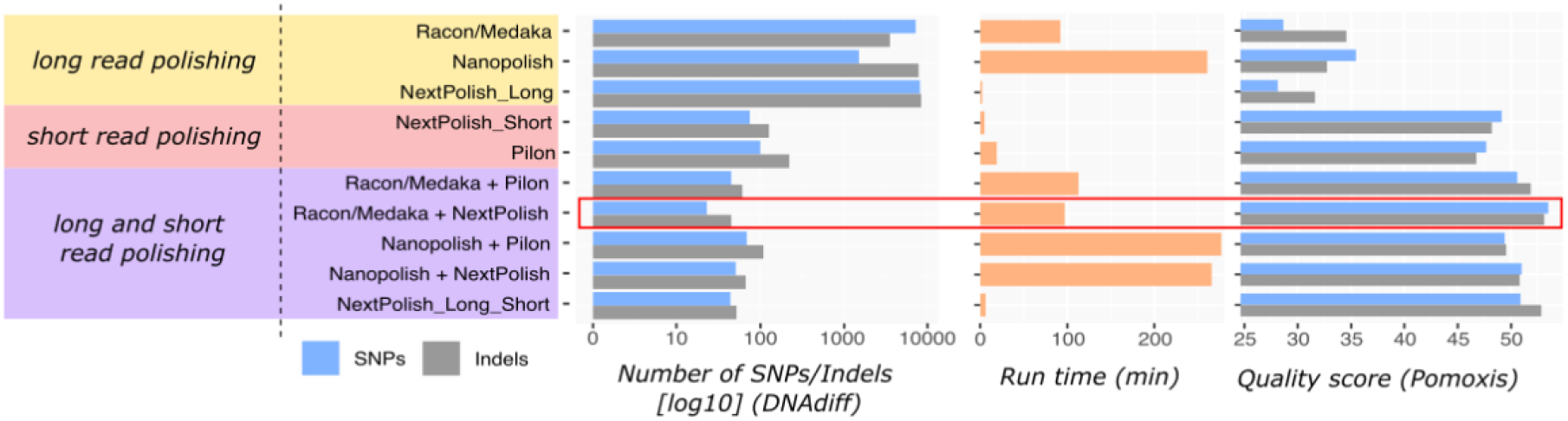
Polishing results for EC958 ONT Flye assembly: Comparative analysis of (i) long read polishing only, (ii) short read polishing only, and (iii) sequential long read and short read polishing, using various tool combinations. Comparison metrics were the number of SNPs/indels to the EC958 reference genome standard (by DNAdiff), run time and quality score (by Poxomis assess_assembly).

To confirm our choice of Flye as the best assembler, we polished assemblies generated from the other five long-read assemblers, described above, using this strategy (**Supplementary Table 5**). The polished Flye assembly remained the most accurate, closely followed by the polished Raven assembly.

#### Hybrid assembly

In addition to long-read assembly (followed by short-read polishing), hybrid assemblers capable of using both long and short reads simultaneously have also been developed, and include Unicycler, MaSuRCA and SPAdes. Comparison of these pipelines to our genome completed with Flye, Racon, Medaka and NextPolish found that they did not outperform our current method. Unicycler was the only hybrid assembler able to completely resolve the chromosome and both plasmids (SPAdes failed to circularise the chromosome while MaSuRCA was unable to assemble the 4 kb plasmid) (**Supplementary Table 6**). Additional long and short read polishing greatly improved the accuracy of the Unicycler and SPAdes hybrid assemblies but not MaSuRCA (**Supplementary Table 5**). We compared the quality of the genomes generated by either the best long-read only assembly (Flye) or the best hybrid assembler based on accuracy and structure (Unicycler) and polished with the same strategy. The polished assemblies contained a similar number of indels compared to the EC958 reference genome standard, however the Flye assembly contained around two-fold fewer substitution errors (**Supplementary Table 5**). Furthermore, Flye was nearly eight times faster than Unicycler (**Supplementary Table 6)**.

### Overall pipeline

Based on the results of our comparative analysis for all of the major steps of bacterial genome assembly, we have developed MicroPIPE **(Figure 4)**. The pipeline is written in Nextflow [20] and the dependencies are packaged into Singularity [21] container images available through the Docker Hub and Quay.io BioContainers repositories. The bioinformatics workflow manager Nextflow allows users to run the pipeline locally or using common High-Performance Computing schedulers. Each step of the pipeline uses a specific container image which enables easy modifications to be made in the future to include new or updated tools. The pipeline is freely available on Github: https://github.com/BeatsonLab-MicrobialGenomics/micropipe.

**Figure 4:**
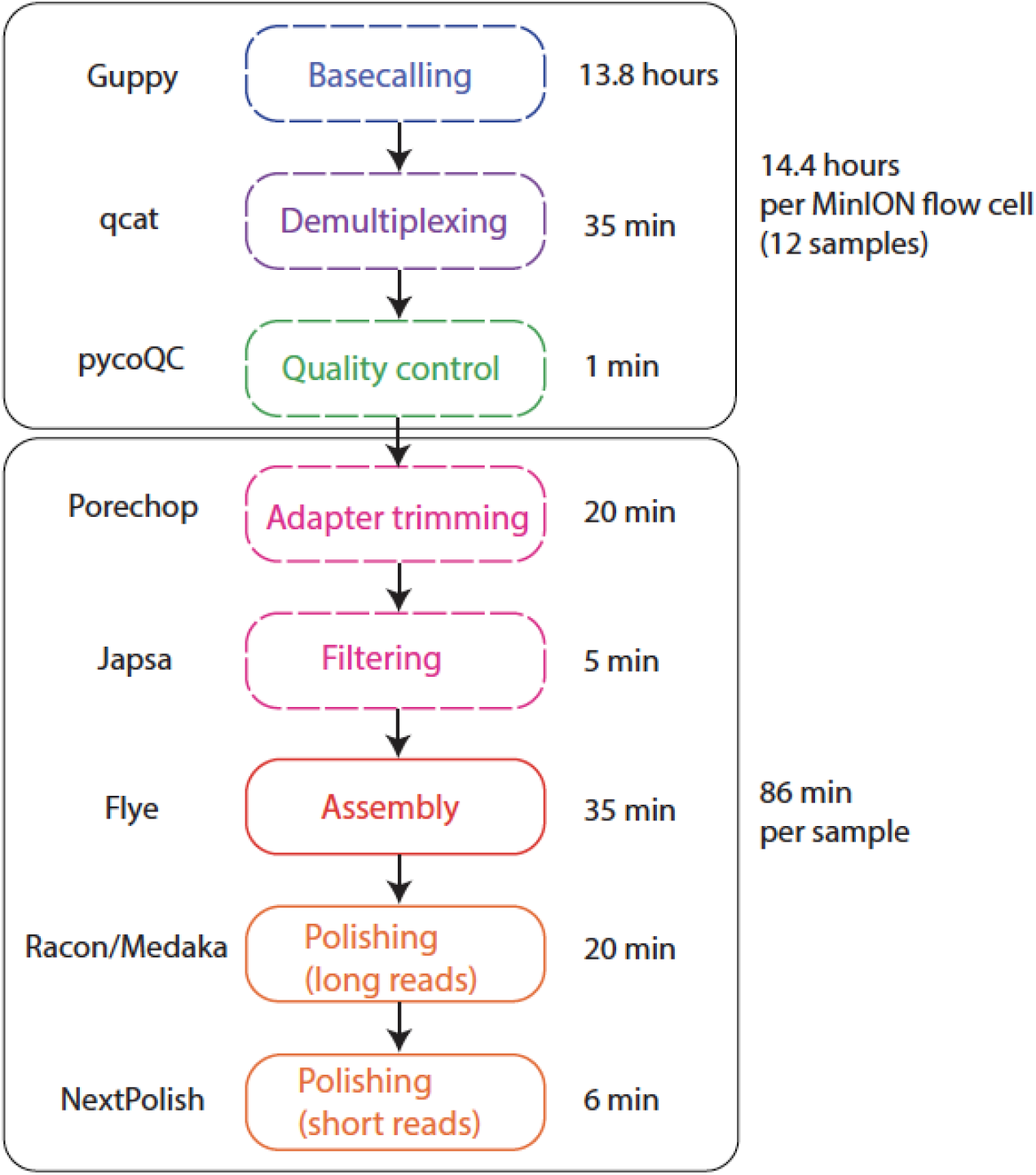
Overall pipeline: Steps involved in genome assembly and the default tool selected for each stage. Steps with dotted outline are optional. Time for running each step is provided based on running 12 multiplexed *E. coli* samples with MicroPIPE v0.8. Basecalling (Guppy) and long-read polishing (Racon and Medaka) were run on a GPU node. The rest of the pipeline was run using CPU resources.

### Evaluation of remaining differences with EC958 reference genome standard

The final genome for EC958 produced by MicroPIPE v0.8 was compared to the previously published EC958 reference genome standard (GenBank: HG941718.1) to assess any remaining differences. We observed a single 3.4 kb inversion corresponding to a phage tail protein switching event previously characterised in EC958 [19]. Overall, there were no other structural rearrangements. MicroPIPE assembled an additional ~1.8 kb plasmid, with 100% nucleotide identity to previously reported *E. coli* plasmids (GenBank records CP048320.1, KJ484633.1, [22]). This plasmid appears to have been lost during size selection when constructing the original genomic DNA library for PacBio RSII sequencing of EC958 as it could be identified from *de novo* assembly of the corresponding Illumina reads.

Comparison of the two assemblies identified 68 remaining differences (66 on the chromosome, 2 on pEC958) (for full list, please see **Supplementary Dataset 1**). The two differences in the plasmid sequence correspond to known errors in the EC958 reference genome standard (PacBio assembly constructed without Illumina polishing). The majority of the chromosomal differences were indels (n=45, 67%) ranging from 1-6 bp in size. These indels were mainly found in rRNA (n=31), tRNA (n=4), insertion sequences (n=4), or phage-related genes (n=2). The remaining 23 differences were SNPs, which were similarly found mainly in rRNA (n=13) and insertion sequences (n=8). These remaining differences likely represent an inability of current short-read polishing to adequately determine true alleles in repetitive regions of the genome. Using methylation-aware basecalling was found to significantly improve these errors, with only 3 SNPs and 31 indels (**Supplementary Table 7**).

During preparation of this manuscript, Guppy v3.6.1 was released. MicroPIPE v0.9 (Guppy v3.6.1) was able to resolve 21 out of the 23 SNPs and 32 out of 45 indels compared to the MicroPIPE v0.8 assembly (Guppy v3.4.3) **(Supplementary Dataset 1, Supplementary Figure 3, Supplementary Table 7)**, relative to the published genome. Two SNPs and 12 indels were additionally detected using v3.6.1, which were not detected using v3.4.3. Both SNPs were detected in IS elements, while 11 out of the 12 indels were detected in rRNA genes. Overall, the v3.6.1 assembly performed better than the v3.4.3 assembly with only 29 differences compared to the complete EC958 genome (4 SNPs and 25 indels). Interestingly, using methylation-aware basecalling with Guppy v3.6.1 was not found to improve overall assembly accuracy **(Supplementary Table 7)**.

### Section 2: Validation of 11 ST131 *E. coli*

To further test the robustness of MicroPIPE on other genomes, we included an additional 11 well-characterised ST131 *E. coli* strains [16] on a multiplexed run of 12 *E. coli* (in addition to EC958).

Each strain took on average 86 minutes to run completely through MicroPIPE v0.8 using 16 threads (excluding the basecalling and demultiplexing steps) **(Figure 4)**. Of these 11 isolates, all had complete circularised chromosomes of the expected size. They also carried an array of plasmids, which were circularised in all cases except for a single isolate, HVM2044 (**Supplementary Table 8**). Re-analysis of this sample found that complete circularised plasmids can be achieved by adjusting the read filtering step. We also identified additional small plasmids in seven out of the 12 genomes ranging between 1.5-5 kb in size. Importantly, we found that these plasmids are not recovered when using filtering parameters above 1 kb.

In order to confirm the accuracy of the assemblies generated with MicroPIPE, we recreated the ST131 phylogeny from [16] using (i) the complete MicroPIPE assembly, (ii) long read only polished assembly, (iii) short read only polished assembly and (iv) unpolished Nanopore assembly, and assessed the position of each strain within the tree. We found that all MicroPIPE v0.8 assemblies and ONT assemblies polished with Illumina clustered closest to their Illumina counterpart within the phylogenetic tree **(Figure 5A)**. However, the long read polished and unpolished ONT assemblies in most cases did not cluster as expected. They also displayed longer branches indicative of the remaining errors within the assembly. Interestingly, the long read polished and unpolished assemblies for all ST131 isolates belonging to our previously defined fluoroquinolone-resistance clade C [16, 17] clustered together independent of other clade C strains, possibly representing systematic errors from the ONT data. Further interrogation of the branch leading to this cluster identified 401 shared SNPs. Of these SNPs, 97% were transitions, particularly A -> G (n=187) and T -> C (n=203) **(Supplementary Table 9, Figure 5B)**. Further analysis of these sites determined that 393 (98%) were associated with a Dcm methylase motif CC(A/T)GG **(Supplementary Figure 4)**.

**Figure 5:**
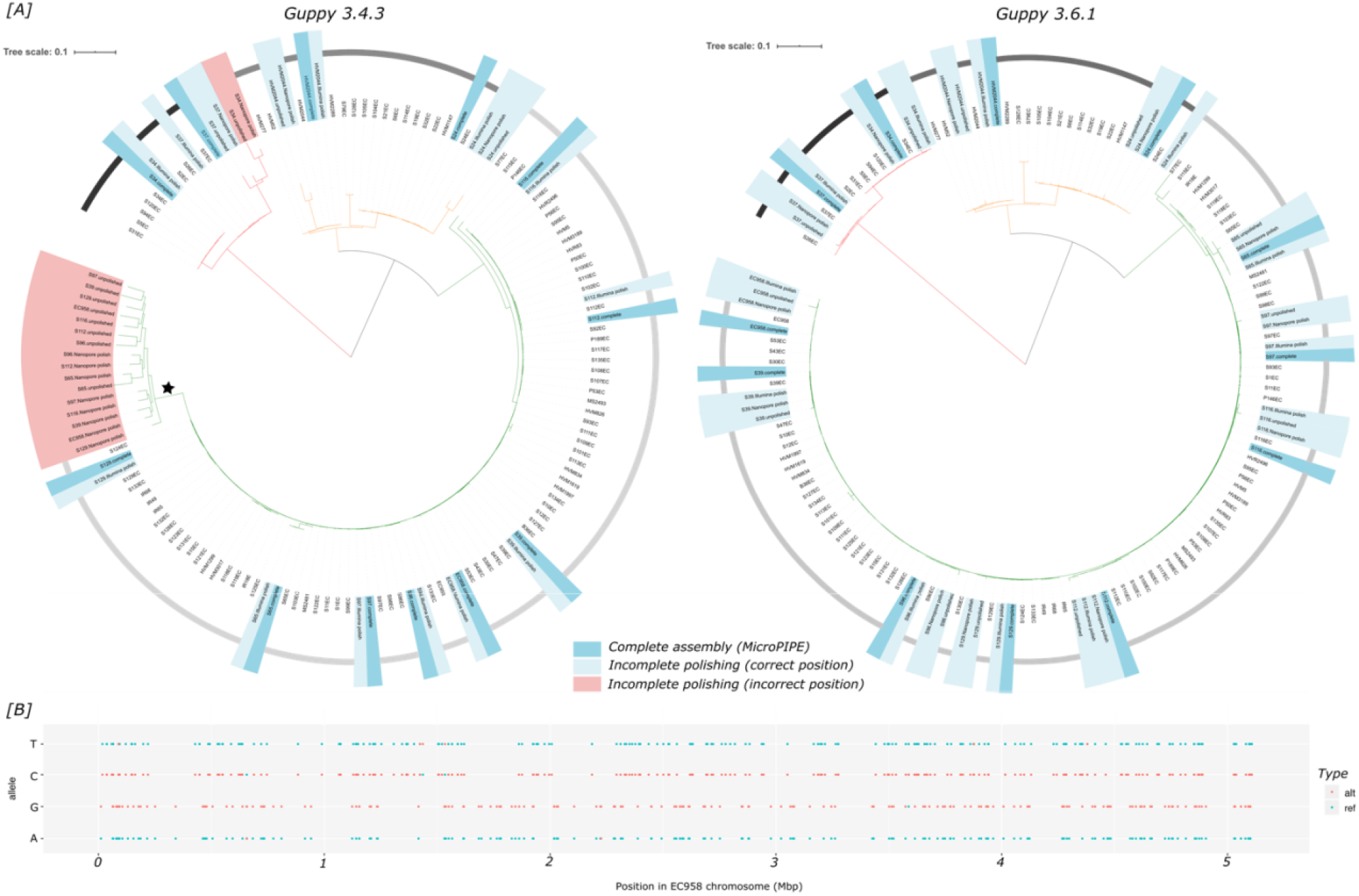
ST131 Phylogeny to assess quality of ONT assemblies: **[A]***dark blue*: Complete polished assemblies from the MicroPIPE pipeline next to their Illumina assembly counterpart in the tree, *light blue*: assemblies with incomplete polishing (i.e. Illumina only, Nanopore only or no polishing) clustered with their Illumina counterpart, *red*: discrepant clustering of Nanopore assemblies. [B] position of alt alleles compared to the EC958 reference standard chromosome present on branch leading to discrepant ONT assemblies as indicated by the star in (A).

We also evaluated the MicroPIPE v0.9 assemblies using Guppy v3.6.1, which was released during preparation of this manuscript. By re-basecalling and recreating all assemblies as before, we found a remarkable increase in the accuracy of Nanopore-only assemblies, such that all assemblies clustered in their expected position within the tree **(Figure 5A)**.

### Section 3: MicroPIPE validation using publicly available ONT sequenced bacteria

Lastly, we tested MicroPIPE using eight public genomes from both Gram-positive and Gram-negative bacteria with available raw nanopore data (fast5) and validated our results using their corresponding publicly available complete genomes (**Supplementary Dataset 1, Supplementary Table 10**). As most of these isolates were sequenced using entire flow cells, the coverage was reduced to 100x during the initial Flye assembly stage to minimise processing time.

Using MicroPIPE v0.9, we were able to completely assemble the chromosome and plasmids of all eight isolates. We were also able to recover two additional plasmids from the *Salmonella enterica* str. SA20055162 that were not reported in the original assembly (**Supplementary Table 10**).

To determine the accuracy of MicroPIPE, we compared our final assemblies with the submitted complete genome for each isolate. Overall, the fewest differences were detected between our MicroPIPE assembly and the complete genome of *Streptococcus pyogenes* strain SP1336, constructed using PacBio long-read sequencing (8 SNPs, 96 indels; **Supplementary Table 10**). All other comparisons yielded 25-510 SNPs, and 14-758 indels, with the worst overall being the *Salmonella enterica* serovar Napoli strain LC0541/17 (**Supplementary Table 10**).

With the exception of *S. pyogenes* SP1336, all other complete genomes were constructed using previously assembled nanopore data (**Supplementary Dataset 1**). As such, we hypothesise that our MicroPIPE assemblies likely represent corrections to the existing complete genomes, as a result of updated basecalling and assembly methods. Further investigation found that one sample, *Salmonella enterica* Bareilly str. CFSAN000189, also had a corresponding complete genome constructed using PacBio data. Comparison of our MicroPIPE assembly to this complete genome detected 0 SNPs and 15 indels, while there were 32 SNPs and 34 indels compared to the ONT complete genome.

## Discussion

ONT long-read sequencing has quickly become one of the most prominent sequencing platforms for microbial researchers globally. However, despite the large number of bacterial genomes being completed using ONT, few end-to-end genome assembly pipelines exist. Here we created an easy, automated and reproducible genome assembly pipeline for the construction of complete, high-quality genomes using ONT in combination with Illumina sequencing. We also provide a robust, publicly available set of 12 ST131 genomes that can be used to validate future pipeline development or software advancements.

One of the main benefits of nanopore sequencing is its cost effectiveness, particularly when multiplexing several samples onto a single flow cell. Methods have been developed to improve yield and length during DNA extraction in order to achieve longer sequencing reads [14, 23].

However, here we show with our method that high-quality complete genomes can be achieved using a standard, commercially available DNA extraction kit coupled with up to 12 multiplexed samples. This build on other advances such as those described by Wick *et al.* [24], and establishes an updated packaged pipeline that provides an efficient, cost effective and reproducible approach to bacterial genome construction.

In our comparative analysis of different aspects of bacterial genome assembly, we chose not to explore the effect of basecallers outside of ONT Guppy basecaller. This comparison has already been completed previously [13], where it was found that Guppy outperformed other existing basecallers. Guppy is also the default basecaller coupled with several of Oxford Nanopore’s devices, such as the MinIT, PromethION and GridION. For these reasons, we felt that it was in the best interest of the community to provide a pipeline that used Guppy as the basecaller. We also made a point of testing both the “high accuracy” mode on a GPU server compared to the “fast” mode on a CPU server, as not all Nanopore users would have access to GPU facilities. We found that, while the GPU server was significantly faster, basecalling reads using the “fast” mode with CPUs can also achieve high-quality genomes with MicroPIPE.

During preparation of this manuscript, Guppy v3.6.1 was released with a raw read accuracy of >97% using R9.4.1 flow cells (https://nanoporetech.com/accuracy). Community feedback regarding this upgraded version supported increased overall accuracy, which prompted us to incorporate this version into our analysis (MicroPipe v0.9). We also found that Guppy v3.6.1 increased the overall accuracy of our assemblies, particularly where it came to unresolved indels using v3.4.3, which were suspected to be the result of technical artefacts around methylated sites [23]. Using Guppy v3.6.1 made Nanopore-only assemblies more feasible, particularly in cases where sufficient genetic context can be provided (e.g. identification of outbreak vs. non-outbreak strains). However, we found that overall both v3.4.3 and v3.6.1 still required polishing with short-read Illumina for maximum accuracy.

We observed some redundancy in the choice of tools for demultiplexing. Binning of reads with both Guppy_barcoder and qcat performed almost equivalently (in terms of number of reads binned), with minimal differences in the overall assembly **(Supplementary Table 11)**. Recent improvements to Guppy_barcoder, which were released by ONT after compilation of this manuscript, suggest that Guppy_barcoder is likely to be the default standard moving forward. MicroPIPE implements a modest filtering measure to remove shorter, low quality reads from the dataset. In this study, we found that filtering reads below 5 kb had little effect on the final chromosome and larger plasmids, while filtering above 1 kb resulted in the loss of several small plasmids in a number of strains (**Supplementary Tables 12 and 13**). Filtering with Filtlong at “--min-length 1000 --keep_percent 90” resulted in the loss of the additional ~1.8 kb small plasmid identified in EC958, which was retained when filtering with Japsa at “--min-length 1000”. As such, we have implemented a 1 kb filtering cut-off (using Japsa) as default in MicroPIPE to retain reads and small plasmids. However, we also found when testing MicroPIPE on publicly available data that harsher filtering is sometimes desirable, especially in cases where a single bacterial genome has been sequenced using an entire flow cell (such that we used the Flye parameter “--asm-coverage 100” to reduce coverage for initial disjointig assembly). As such, pre-processing of large quantities or highly ununiform data using Filtlong may be the most desirable method. Ultimately, understanding the quality and read lengths of the input data is a valuable step in generating the best possible assembly. We also provided the user read quality assessment using PycoQC to assist in parameter selection.

Several other comparative analyses have been published exploring the overall utility of different assemblers, in particular Wick *et al.* [25], who provide a comprehensive assembly comparison using both simulated and real read datasets. While we did not test NECAT and Miniasm, we found that our results generally matched those reported by Wick *et al*., particularly when it came to the overall strong performance of Flye. The most recent version of Flye (v2.8) also removes the need to nominate a genome size, making it a more robust option. However, we found that this version did not outperform the release used in this paper (v2.5) on our dataset, as it was unable to circularise all plasmids. As such, we have retained Flye v2.5 in MicroPIPE.

Long and short read polishing is a staple of high-quality genome assembly, as the combination of both ensures the correct contextual placement of variants as well as highly accurate basecalls. However, while long-reads have enabled completion of assemblies by spanning repetitive regions, polishing of these regions with short reads remains a problem. Here we found that the majority of remaining differences between our EC958 ONT assembly and the reference assembly (constructed with PacBio single molecule real time [SMRT] sequencing) resided in repetitive regions. Ideally, polishing with long reads only would be a viable method to reduce these errors as they would have sufficient coverage to ensure correct placement of the repeat variant. However, as we show here, long read-only polishing was insufficient (likely due to per-read accuracy), and short read polishing was necessary for removal of the majority of errors. Currently, final polishing and assembly prior to completion will still necessitate manual frameshift inspection. While impractical and costly, a combination of both PacBio and ONT assembly could correct inherent biases in both technologies, using a consensus tool such as Trycycler (https://github.com/rrwick/Trycycler). Long-read correction could also provide another means of error reduction, however, was not assessed in this paper [26, 27].

We validated MicroPIPE using a set of 12 well-characterised *E. coli* isolates described previously from a global collection [16, 17]. We did this for several reasons, including (i) the availability of an existing high-quality reference genome and associated phylogenetic data (ii) the robustness of *E. coli* as a representative species and workhorse organism, and (iii) our extensive knowledge of the *E. coli* genome and ST131 lineage. We hope that by providing this dataset to the wider community, it can serve as a resource for future validation and testing of not only MicroPIPE, but other microbial assembly pipelines and tools.

In addition to in-house ONT sequencing data, we also tested MicroPIPE on a variety of publicly available bacterial genomes to evaluate its assembly capabilities on other species. Without any manual intervention, MicroPIPE was able to assemble all eight genomes, while also recovering additional plasmids that were likely missed in the original assembly. When evaluating correctness of the genomes, we found a number of remaining SNPs and indels when compared to the complete genomes provided. Investigation into construction of the reference genomes found that seven of the eight genomes provided were constructed previously using ONT sequencing data, leading us to believe that differences in our assemblies compared to the “reference” genomes may actually be corrections. Indeed, the genome with the closest match between reference and MicroPIPE assembly were the genomes constructed using PacBio. As such, we believe that genomes completed historically using ONT reads should be used cautiously, and raw ONT data provided where possible to allow for reconstruction and improvement of the assembly as the technology improves.

## Conclusions

Overall, we present an end-to-end pipeline for high-quality bacterial genome construction designed to be easily implemented in the research lab setting. We believe this will be a useful resource for users to easily and reproducibly construct complete bacterial genomes from Nanopore sequencing data.

## Methods

### Public data

The EC958 complete genome was downloaded from NCBI (GenBank: HG941718.1, HG941719.1, HG941720.1) [19]. Illumina reads for 12 ST131 genomes and draft assemblies for 95 ST131 were accessed from [16]. Eight publicly available complete genomes were also selected to test MicroPIPE, under the following criteria: (i) the raw nanopore sequencing files (fast5) were available, (ii) a complete genome was made available for the same strain and (iii) Illumina sequencing data were available for the same strain. These eight genomes represented 5 species from both gram-positive and gram-negative bacteria with chromosome sizes between 1.8 Mbp – 5.5 Mbps. A complete list of data used is provided in **Supplementary dataset 1**.

### Culture and DNA extraction

12 ST131 *E. coli* isolates (including EC958) were grown from single colonies in Lysogeny Broth (LB) at 37°C overnight with 250 rpm shaking. The overnight cultures (1.5 mL) were then pelleted for DNA extraction using the Wizard Genomic DNA Purification Kit (Promega) following manufacturer’s protocol with modifications. Briefly, the cell pellet was lysed following the protocol for Gram negative bacteria. RNA was removed by 1h incubation at 37°C with RNase and the lysate was then mix with Protein Precipitation Solution by vortexing for 5s at max speed using Vortex-Genie 2 with horizontal tube adapter (Scientific Industries). The DNA was precipitated using isopropanol and washed with 70% ethanol. The DNA pellet was air-dried and then rehydrated in 100 μl EB buffer (QIAgen) by incubation at 65°C for 1 hour. The DNA was quantified using a Qubit fluorometer (ThermoFisher Scientific) and the DNA fragment size was estimated using agarose gel electrophoresis (0.5% agarose in TAE, 90V, 1h30m).

### Nanopore sequencing

DNA from 12 ST131 *E. coli* were multiplexed onto a single FLO-MIN106 flow cell using the rapid barcode sequencing kit (SQK-RBK004) as per manufacturer’s recommendation with the following adjustments: the barcoded DNA was pooled without a concentration step using AMPure XP beads prior to sequencing. Read metrics for each isolate are given in **Supplementary table 1**.

### Pipeline tools and settings

Specific parameters and commands used to perform the following analyses are provided in full in **Supplementary dataset 1**. MicroPIPE v0.8 uses Guppy v3.4.3, while MicroPIPE v0.9 uses Guppy v3.6.1.

#### Basecalling

Reads were basecalled using Guppy (v3.4.3) “fast” and “high-accuracy” modes. Fast mode was evaluated using both GPU and CPU servers, while the “high-accuracy” mode was evaluated using only GPU as the time to completion for this mode became unfeasible when run using CPUs. Upon the release of Guppy v3.6.1, reads were re-basecalled using only the “high-accuracy” mode. Guppy versions (3.4.3 and 3.6.1) were tested using the methylation aware config file “dna_r9.4.1_450bps_modbases_dam-dcm-cpg_hac.cfg”.

#### Demultiplexing

Demultiplexing was evaluated using Guppy_barcoder (v3.4.3) and qcat (v1.0.1) on the “passed” (>Q7) fastq reads after basecalling with Guppy. Demultiplexing using the raw fast5 reads was evaluated using Deepbinner (v0.2.0) [28]. Demultiplexed fast5 reads were subsequently basecalled with Guppy (v3.4.3).

#### Quality control

Barcodes and adapters were trimmed using Porechop (v0.2.3_seqan2.1.1) (https://github.com/rrwick/Porechop). Overall read quality metrics and basecalling statistics were extracted using PycoQC (v2.2.3) [29]. Read length and quality metrics per sample were extracted using NanoPlot (v1.26.1) [30]. Average percentage read accuracy was determined by mapping the basecalled reads to the reference genome EC958 using Minimap2 (v2.17-r954-dirty) [31] and computing reads accuracy using Nanoplot. Filtering was evaluated using two tools: Filtlong (v0.2.0) (https://github.com/rrwick/Filtlong) and Japsa (v1.9-01a) (https://github.com/mdcao/japsa/).

#### Assembly

Six assemblers were evaluated for long-read assembly only: Canu (v1.9) [32], Flye (v2.5) [33], Raven (v1.1.5) (https://github.com/lbcb-sci/raven), Redbean (v2.5) [34], Shasta (v0.4.0: config file optimised for Nanopore: https://github.com/chanzuckerberg/shasta/blob/master/conf/Nanopore-Dec2019.conf) [35] and Unicycler (v0.4.7 long-read only) [36]. Three hybrid-assembly tools were also evaluated, including SPAdes (v3.13.1) [37], Unicycler (v0.4.7) and MaSuRCA (v3.3.5) [38].

#### Polishing and quality assessment

Polishing of the draft assemblies was evaluated using long reads (ONT), short reads (Illumina), and a combination of both long and short reads. Long read polishing was performed using Racon (v1.4.9) [39] and Medaka (v0.10.0) (https://nanoporetech.github.io/medaka/) (4 iterations of Racon based on Minimap2 v2.17-r941 overlaps followed by one iteration of Medaka), Nanopolish (v0.11.1) [40] (1 iteration based on Minimap2 v2.17-r941 alignment) and NextPolish (v1.1.0) [41] (2 iterations). Raw Illumina reads were trimmed using Trimmomatic (v0.36) [42] with the following settings: ILLUMINACLIP:TruSeq3-PE-2.fa:2:30:10 SLIDINGWINDOW:4:20 MINLEN:30. Short read polishing was performed using NextPolish (v1.1.0) and Pilon (v1.23) [43] (both 2 rounds of polishing based on BWA MEM v0.7.17-r1188 alignments).

Circularity was checked using NUCmer (v3.1) [44] to perform self-alignments. Final assemblies were assessed for quality by comparison to the complete EC958 genome using the assess_assembly tool from Pomoxis (v0.3) (https://github.com/nanoporetech/pomoxis) as well as DNAdiff (v1.3) [44] and QUAST (v5.0.2) [45] to detect errors, misassemblies, and determine overall nucleotide identity.

#### Compute resources

All results were produced using cloud-based nodes with 16vCPUs and 32GB RAM. For the GPU node, the GPU is a NVIDIA Tesla P40 24GB while the CPUs are 2x Intel Xeon Silver 4214 2.2G (12C/24T, 9.6GT/s, 16.5M Cache, Turbo, HT [85W] DDR4-2400).

### ST131 phylogeny

Parsnp (v1.5.2) [46] was used to create an ST131 phylogeny using the 12 ST131 *E. coli* assembled in this study in addition to 95 ST131 *E. coli* short-read assemblies from Petty and Ben Zakour *et al.* [16]. Recombination was removed using PhiPack [47], as implemented in Parsnp. To evaluate the accuracy of each assembly and polishing step, we included our 12 completely polished assemblies (long and short read), 12 unpolished assemblies, 12 long-read polished assemblies and 12 short-read polished assemblies. The tree was visualised using Figtree (http://tree.bio.ed.ac.uk/software/figtree/) and iTOL [48].

### MEME methylation motif analysis

The 20 bps sequence (−10 to +10) around the 401 shared SNPs were extracted using BEDTools getfasta (v2.28.0-33-g0f45761e) [49]. MEME (v5.2.0) [50, 51] was used to identify enriched motifs within the sequences using the default parameters of the classic mode and allowing zero or one occurrence per sequence. The motif CC(T/A)GG was significantly enriched in 393 sequences with an E-value of 6.2e-758.

## Declarations

### Ethics approval and consent to participate

Not applicable.

### Consent for publication

Not applicable.

### Availability of data and materials

The datasets generated and analysed during the current study are available under the following Bioprojects (specific accessions available in supplementary dataset 1): EC958 complete genome (GenBank: HG941718.1), ST131 Illumina data (PRJEB2968), ST131 Nanopore data (fast5 and fastq [demultiplexed]; PRJNA679678).

### Competing interests

None to declare.

### Funding

LWR was supported by a Sakzewski Translational Research Grant. This work was supported by funding from the Queensland Genomics Health Alliance (now Queensland Genomics), Queensland Health, the Queensland Government.

### Authors’ contributions

All authors conceptualised the study. VM, LWR, BMF and SAB developed the methodology. MDP and MAS provided the bacterial strains and ONT sequencing data. VM wrote the pipeline. VM, LWR and NTKN conducted formal analysis. All authors contributed to the interpretation of results. SAB and MAS supervised aspects of the project and provided essential expert analysis. LWR and VM wrote the original manuscript. BMF and SAB edited the manuscript. All authors read and approved the final manuscript.

## Acknowledgements

We would like to acknowledge Thom Cuddihy (QCIF Facility for Advanced Bioinformatics) for his assistance and advice regarding pipeline testing and development. This research was supported by QRIScloud and by use of the Nectar Research Cloud. The Nectar Research Cloud is a collaborative Australian research platform supported by the National Collaborative Research Infrastructure Strategy (NCRIS).

## Notes

### Competing Interest Statement

The authors have declared no competing interest.

