## Supplementary file for "MicroPIPE: *An end-to-end solution for high-quality complete bacterial genome construction*"

### Supplementary Materials

#### Supplementary Table 1: Read length and quality metrics per isolate

| Sample | Number of Reads | Number of Bases | Read Length N50 | Median Read Length | Median Read Quality |
|---|---|---|---|---|---|
| S24 | 83,023 | 518,350,475 | 11,366 | 3,870 | 11.7 |
| S34 | 130,069 | 998,292,413 | 13,795 | 4,900 | 11.6 |
| S37 | 82,516 | 524,209,315 | 11,692 | 3,938 | 11.6 |
| S39 | 127,098 | 1,021,807,859 | 14,852 | 4,933 | 11.6 |
| S65 | 93,069 | 701,752,024 | 13,675 | 4,742 | 11.7 |
| S96 | 138,299 | 901,024,145 | 11,778 | 4,111 | 11.6 |
| S97 | 149,425 | 1,138,211,010 | 13,728 | 4,820 | 11.7 |
| S112 | 91,655 | 705,206,951 | 14,213 | 4,740 | 11.6 |
| S116 | 111,062 | 900,213,042 | 14,681 | 5,131 | 11.6 |
| S129 | 66,930 | 505,613,353 | 14,242 | 4,470 | 11.6 |
| EC958 | 242,401 | 1,755,197,084 | 13,131 | 4,577 | 11.6 |
| HVM2044 | 117,328 | 778,919,514 | 12,001 | 4,145 | 11.5 |

**Supplementary Table 2: assembly results for EC958**

| Assembler | Runtime (min) | Number of contigs | Size (bp) | Nucleotide Identity (%) | Number of misassemblies and local misassemblies | Genome fraction (%) | Circular? |
|---|---|---|---|---|---|---|---|
| Flye v2.5 | 46 | 4 | 5,130,397<br>136,261<br>4,243<br>1,841 | 99.41 | 2<br>1 | 100 | Yes<br>Yes<br>Yes<br>Yes |
| Canu v1.9 | 168 | 5 | 5,091,050 *<br>135,078 *<br>4,078 *<br>1,814 *<br>1,898 | 99.60 | 4<br>4 | 99.873 | Yes<br>Yes<br>Yes<br>Yes (nucmer)<br>No |
| Unicycler (long-read) v0.4.7 | 83 | 6 | 5,104,112<br>133,747<br>111,708<br>107,098<br>57,320<br>4,085 | 99.71 | 13<br>3 | 99.961 | Yes<br>Yes<br>No<br>No<br>No<br>Yes |
| Raven v1.1.5 | 19 | 2 | 5,103,450<br>135,356 | 99.73 | 0<br>1 | 99.863 | Yes<br>Yes |
| Redbean v2.5 | 8.5 | 2 | 5,089,255<br>135,180 | 99.59 | 2<br>2 | 99.831 | Yes (manual)<br>Yes (manual) |
| Shasta v0.4.0 | 4.5 | 2 | 5,113,702<br>135,731 | 99.31 | 0<br>9 | 99.921 | Yes<br>Yes |

* : Sizes correspond to contigs after manual trimming of overlapping ends

**Supplementary Table 3: EC958 assembly results using different Flye parameters, demultiplexing tools and read filtering parameters:** This table summarised the number of contigs from the Flye assembly and the number of contigs reported as circular using different read filtering parameters, different demultiplexing tools and different Flye parameters. The numbers represent “number of circular contigs” : “total number of contigs assembled”. *Note:* the expected number of contigs for the complete EC958 genome is 4 (1 chromosome, and 3 plasmids). Any assemblies with less than 4 total contigs were found to have plasmids that were not assembled.

| **Filtering** | **Demultiplexing** | **Number of reads** | **Number of circular contigs : Number of contigs in assembly** | | |
|---|---|---|---|---|---|
| | | | **Flye (default)** | **Flye --plasmids** | **Flye --plasmids --meta** |
| All reads (Porechop trimmed) | Deepbinner | 182,607 | 2:2 | 4:4 | 4:4 |
| | Guppy | 226,464 | 2:2 | 4:4 | 4:4 |
| | Qcat | 238,000 | 2:2 | 4:4 | 4:4 |
| Japsa | Deepbinner | 153,573 | 2:2 | 4:4 | 4:4 |
| --lenMin 1000 | Guppy | 190,440 | 2:2 | 4:4 | 4:4 |
| --qualMin 10 | Qcat | 200,148 | 2:2 | 4:4 | 4:4 |
| Japsa | Deepbinner | 131,623 | 2:2 | 3:3 | 3:3 |
| --lenMin 2000 | Guppy | 163,137 | 2:2 | 3:3 | 3:3 |
| --qualMin 5 | Qcat | 171,456 | 2:2 | 3:3 | 3:3 |
| Filtlong | Deepbinner | 104,487 | 2:2 | 3:3 | 3:3 |
| --min_length 1000 | Guppy | 129,170 | 2:2 | 3:3 | 3:3 |
| --keep_percent 90 | Qcat | 135,791 | 2:2 | 3:3 | 3:3 |

**Supplementary Table 4: Polishing tool comparison:** Racon/Medaka + NextPolish (green) was selected for MicroPIPE

| | Read set | Run time (min) | DNAdiff | | | Pomoxis | | | QUAST | |
|---|---|---|---|---|---|---|---|---|---|---|
| | | | Nucleotide Identity (%) | SNPs | Indels | Quality score | Identity quality score | Indel quality score | Mismatches per 100 kb | Indels per 100 kb |
| Racon/Medaka | Long | 92 | 99.79 | 7199 | 3536 | 26.85 | 28.65 | 34.56 | 136.57 | 67.66 |
| Racon/Medaka + Pilon | Long and short | 113 | 99.99 | 45 | 61 | 46.34 | 50.6 | 51.86 | 0.86 | 1.18 |
| Racon/Medaka + NextPolish | Long and short | 97 | 99.99 | 23 | 45 | 48.10 | 53.52 | 53.14 | 0.44 | 0.88 |
| Nanopolish | Long | 261 | 99.82 | 1512 | 7807 | 27.43 | 35.46 | 32.74 | 28.35 | 148.83 |
| Nanopolish + Pilon | Long and short | 277 | 99.99 | 69 | 109 | 44.35 | 49.42 | 49.55 | 1.07 | 2.1 |
| Nanopolish + NextPolish | Long and short | 266 | 99.99 | 51 | 67 | 45.93 | 51.01 | 50.82 | 0.72 | 1.3 |
| NextPolish | Long | 2.5 | 99.67 | 8108 | 8367 | 24.81 | 28.13 | 31.61 | 153.43 | 159.94 |
| NextPolish | Short | 5 | 99.99 | 75 | 127 | 43.26 | 49.14 | 48.2 | 1.22 | 2.42 |
| NextPolish | Long and short | 6.5 | 99.99 | 44 | 52 | 46.92 | 50.9 | 52.85 | 0.8 | 1.03 |
| Pilon | Short | 19 | 99.99 | 100 | 222 | 41.42 | 47.69 | 46.75 | 1.7 | 4.23 |

**Supplementary Table 5: Polishing comparison using other assemblers (Long read assembly followed by polishing with Racon/Medaka+NextPolish [blue] and hybrid assembly [yellow]):** top three in each quality category highlighted green

| Assembler | Assembly strategy | DNAdiff | | | Pomoxis | | | QUAST | | | | |
|---|---|---|---|---|---|---|---|---|---|---|---|---|
| | | Nucleotide Identity (%) | SNPs | Indels | Quality score | Identity quality score | Indel quality score | Mismatches per 100 kb | Indels per 100 kb | Indels length | Genome fraction (%) | Duplication Ratio |
| Flye v2.5 | Long read assembly + polishing | 99.99 | 23 | 45 | 48.10 | 53.52 | 53.14 | 0.44 | 0.88 | 58 | 100 | 1 |
| Canu v1.9 | Long read assembly + polishing | 99.99 | 51 | 41 | 45.23 | 50.35 | 50.36 | 2.48 | 2.5 | 185 | 99.952 | 1.021 |
| Unicycler v0.4.7 | Long read assembly + polishing | 99.99 | 78 | 65 | 45.54 | 48.58 | 52.01 | 16.26 | 18.47 | 1469 | 99.949 | 1.049 |
| Raven v1.1.5 | Long read assembly + polishing | 99.99 | 28 | 49 | 48.25 | 52.7 | 53.88 | 0.53 | 0.93 | 55 | 99.922 | 1 |
| Redbean v2.5 | Long read assembly + polishing | 99.99 | 28 | 56 | 45.74 | 50.64 | 51.93 | 0.92 | 1.01 | 184 | 99.831 | 1 |
| Shasta v0.4.0 | Long read assembly + polishing | 99.99 | 91 | 38 | 46.88 | 49.89 | 53.68 | 1.66 | 0.82 | 79 | 99.922 | 1 |
| Unicycler v0.4.7 | Hybrid assembly | 99.99 | 165 | 34 | 41.38 | 43.06 | 50.37 | 3.11 | 0.61 | 159 | 100 | 1 |
| | Hybrid assembly + polishing | 99.99 | 38 | 45 | 44.10 | 45.68 | 52.54 | 0.88 | 0.91 | 63 | 100 | 1 |
| MaSuRCA v3.3.5 | Hybrid assembly | 99.99 | 67 | 15 | 39.15 | 49.00 | 42.67 | 1.35 | 0.31 | 32 | 99.922 | 1.022 |
| | Hybrid assembly + polishing | 99.98 | 63 | 49 | 40.77 | 43.01 | 47.95 | 4.94 | 5.66 | 412 | 99.917 | 1.021 |
| SPAdes v3.13.1 | Hybrid assembly | 99.98 | 1073 | 87 | 35.31 | 37.03 | 43.66 | 20.92 | 1.83 | 562 | 99.960 | 1 |
| | Hybrid assembly + polishing | 99.99 | 43 | 37 | 48.10 | 51.06 | 54.67 | 0.78 | 0.74 | 95 | 99.962 | 1 |

**Supplementary Table 6: Hybrid assembly comparison to Flye+Racon/medaka+NextPolish:**

Flye + polishing (green) was selected for MicroPIPE

| | Runtime (min) | Number of contigs | Size of contigs (bp) | Circularised? | DNAdiff (nucleotide identity) | Number of misassemblies | Genome fraction (%) |
|---|---|---|---|---|---|---|---|
| Flye v2.5 only assembly | 46 | 4 | 5,130,406<br>136,260<br>4,245<br>1,841 | Yes<br>Yes<br>Yes<br>Yes | 99.41 | 2 | 100 |
| Flye v2.5 + polishing | 143 | 4 | 5,109,793<br>135,596<br>4,208<br>1,823 | Yes<br>Yes<br>Yes<br>Yes | 99.99 | 2 | 100 |
| Unicycler v0.4.7 (hybrid) | 360 | 4 | 5,109,706<br>135,600<br>4,088<br>1,822 | Yes<br>Yes<br>Yes<br>Yes | 99.99 | 4 | 100 |
| MaSuRCA v3.3.5 | 55 | 2 | 5,149,439 (5,109,995*)<br>210,338 (135,599*) | Yes (nucmer)<br>Yes (nucmer) | 99.99 | 2 | 99.922 |
| SPAdes v3.13.1 | 70 | 5 | 2,937,224<br>2,171,161<br>134,802<br>4,088<br>1,822 | No<br>No<br>Yes (manual)<br>Yes (manual)<br>Yes (manual) | 99.98 | 0 | 99.960 |

* Size after manual circularisation with nucmer

**Supplementary Table 7: final assembly comparisons between Guppy versions and methylation-aware basecalling**

| Guppy version | Guppy model | Data used for polishing | Nb SNPs | Nb Indels | Nb Total |
|---|---|---|---|---|---|
| v3.4.3 | hac | ONT | 7,199 | 3,536 | 10,735 |
| | | ONT + Illumina | 23 | 45 | 68 |
| | modbases_hac | ONT | 160 | 1,997 | 2,157 |
| | | ONT + Illumina | 3 | 31 | 34 |
| v3.6.1 | hac | ONT | 28 | 438 | 466 |
| | | ONT + Illumina | 4 | 25 | 29 |
| | modbases_hac | ONT | 117 | 1,663 | 1,780 |
| | | ONT + Illumina | 5 | 27 | 32 |

#### Supplementary Table 8: MicroPIPE v0.8 results for 11 ST131

| Strain | Chromosome/plasmid | Size (bps) | Circularised? |
|---|---|---|---|
| S24EC | Chromosome<br>Plasmid A | 5,061,955<br>114,236 | Yes<br>Yes |
| S34EC | Chromosome<br>Plasmid A<br>Plasmid B | 5,034,986<br>152,718<br>107,794 | Yes<br>Yes<br>Yes |
| S37EC | Chromosome<br>Plasmid A<br>Plasmid B | 4,965,673<br>157,040<br>60,747 | Yes<br>Yes<br>Yes |
| S39EC | Chromosome<br>Plasmid A<br>Plasmid B<br>Plasmid C<br>Plasmid D<br>Plasmid E<br>Plasmid F | 5,038,121<br>143,728<br>94,575<br>66,312<br>61,832<br>2,018<br>1,788 | Yes<br>Yes<br>Yes<br>Yes<br>Yes<br>Yes<br>Yes |
| S65EC | Chromosome<br>Plasmid A | 5,187,864<br>146,795 | Yes<br>Yes |
| S96EC | Chromosome<br>Plasmid A<br>Plasmid B<br>Plasmid C<br>Plasmid D | 5,052,350<br>163,696<br>115,542<br>14,053<br>4,073 | Yes<br>Yes<br>Yes<br>Yes<br>Yes |
| S97EC | Chromosome<br>Plasmid A<br>Plasmid B<br>Plasmid C<br>Plasmid D | 5,162,367<br>165,456<br>96,393<br>4,059<br>3,185 | Yes<br>Yes<br>Yes<br>Yes<br>Yes |
| S112EC | Chromosome<br>Plasmid A<br>Plasmid B<br>Plasmid C<br>Plasmid D | 5,003,915<br>160,386<br>34,185<br>5,208<br>4,141 | Yes<br>Yes<br>Yes<br>Yes<br>Yes |
| S116EC | Chromosome<br>Plasmid A<br>Plasmid B<br>Plasmid C<br>Plasmid D | 4,972,973<br>66,522<br>5,183<br>4,122<br>4,117 | Yes<br>Yes<br>Yes<br>Yes<br>Yes |
| S129EC | Chromosome<br>Plasmid A<br>Plasmid B<br>Plasmid C<br>Plasmid D<br>Plasmid E | 5,177,746<br>163,021<br>93,601<br>33,138<br>4,071<br>2,399 | Yes<br>Yes<br>Yes<br>Yes<br>Yes<br>Yes |

| | | | |
|---|---|---|---|
| | Plasmid F<br>Plasmid G | 2,108<br>1,555 | Yes<br>Yes |
| HVM2044 | Chromosome<br>Plasmid A<br>Plasmid B<br>Plasmid C<br>Plasmid D | 4,986,664<br>142,349<br>115,439<br>18,084<br>6,772 | Yes*<br>No<br>Yes<br>Yes<br>No |

* modifying filtering parameters resulted in 5 circular contigs (Filtlong --min_length 1000 --keep_percent 90)

**Supplementary Table 9: SNP types for clade C unpolished/Illumina unpolished assemblies**

| SNP type | Allele change (ref → alt) | Count |
|---|---|---|
| **Transition**<br>399 | A → G | 187 |
| | G → A | 3 |
| | T → C | 203 |
| | C → T | 6 |
| **Transversion**<br>2 | A → T | 0 |
| | T → A | 0 |
| | T → G | 0 |
| | G → T | 0 |
| | G → C | 1 |
| | C → G | 1 |
| | C → A | 0 |
| | A → C | 0 |

#### Supplementary Table 10: MicroPIPE v0.9 results for public datasets

Flye was run using the --asm-coverage 100 parameter in order to reduce the computational run time. Only circular contigs are reported (as identified by Flye). For further details on all public data, see Supplementary dataset 1.

| Reference | Strain | Reference genome assembly method and coverage | Chromosome/ plasmid | Reference genome size (bps) | Assembly size (bps) | Circular? | Nucleotide Identity (%) | DNAdiff SNPs GSNPs | DNAdiff Indels | QUAST misassemblies |
|---|---|---|---|---|---|---|---|---|---|---|
| Clement et al | Salmonella enterica serovar Napoli strain LC0541/17 | Canu using Nanopore + Illumina 37x | Chromosome<br>pLC0541_17 | 4,679,033<br>90,558 | 4,679,747<br>90,578 | Yes<br>Yes | 99.97 | 510<br>390 | 758 | 0 |
| Sydenham et al | Bacteroides fragilis strain:DCMOUH0042B (BF042) | Unicycler using Nanopore + Illumina 200x | Chromosome<br>pBFO42_1<br>pBFO42_2 | 5,141,257<br>8,306<br>5,594 | 5,141,261<br>8,316<br>5,629 | Yes<br>Yes<br>Yes | 99.99 | 65<br>9 | 14 | 0 |
| Sydenham et al | Bacteroides fragilis strain:CCUG4856T | Unicycler using Nanopore + Illumina 200x | Chromosome<br>pBF9343 | 5,205,133<br>36,560 | 5,205,138<br>36,559 | Yes<br>Yes | 99.99 | 25<br>5 | 22 | 1 (inversion) |
| Walker et al | Streptococcus pyogenes strain SP1336 | Pacbio 105x | Chromosome | 1,878,827 | 1,878,922 | Yes | 99.99 | 8<br>6 | 96 | 0 |
| Wick et al GB | Klebsiella pneumoniae INF032 | Unicycler using Nanopore + Illumina 133x | Chromosome | 5,111,537 | 5,111,663 | Yes | 99.99 | 137<br>72 | 172 | 0 |
| Taylor et al | E. coli O157:H7 strain FSIS11705876 | Unicycler using Nanopore + Illumina 692x | Chromosome<br>pO157 | 5,483,434<br>94,581 | 5,483,452<br>94,593 | Yes<br>Yes | 99.99 | 52<br>2 | 103 | 0 |
| Taylor et al | Salmonella enterica Bareilly str. CFSAN000189 | Unicycler using Nanopore + Illumina 599x | Chromosome<br>Plasmid | 4,724,806<br>81,814 | 4,724,797<br>81,815 | Yes<br>Yes | 99.99 | 32<br>21 | 34 | 0 |
| | | SMRT Analysis v. 1.3.3 using PacBio RS 80x | Chromosome<br>Plasmid | 4,730,612<br>78,193 | | | 99.99 | 0<br>0 | 15 | 3 |
| Bessonov et al | Salmonella enterica str. SA20055162 | Unicycler using Nanopore + Illumina 50x | Chromosome<br>Plasmid<br>Plasmid | 4,640,729 | 4,640,715<br>105,679<br>98,127 | Yes<br>Yes<br>Yes | 99.99 | 53<br>15 | 22 | 0 |

**Supplementary Table 11: Demultiplexing comparison between qcat and Guppy: run-times, EC958 read and assembly accuracy**

| **Basecalling comparison** | **Guppy3.4.3_hac** | | **Guppy3.6.1_hac** | |
|---|---|---|---|---|
| **Demultiplexing tool** | **qcat** | **guppy** | **qcat** | **guppy** |
| Demultiplexing run time (h) | 0.58 | 0.68 | 0.56 | 0.47 |
| Average read percent identity | 91.0 | 91.2 | 93.7 | 93.8 |
| Mean read quality | 11.4 | 11.5 | 13.3 | 13.4 |
| Read length N50 | 13,092 | 13,087 | 13,021 | 13,013 |
| Number of binned reads | 240,766 | 229,100 | 244,830 | 240,124 |
| **Final assembly comparison** | | | | |
| Assembly nucleotide identity (%) | 99.99 | 99.99 | 99.99 | 99.99 |
| Number of SNP (DNAdiff) | 23 | 28 | 4 | 4 |
| Number of GSNP (DNAdiff) | 3 | 6 | 1 | 1 |
| Number of indels (DNAdiff) | 45 | 35 | 25 | 25 |
| Assembly quality score (Pomoxis) | 48.10 | 48.76 | 52.27 | 52.41 |
| Mismatches per 100 kb (QUAST) | 0.44 | 0.53 | 0.08 | 0.08 |
| Indels per 100 kb (QUAST) | 0.88 | 0.69 | 0.50 | 0.48 |
| Contig Size (bp) | 5,109,793<br>135,596<br>4,208<br>1,823 | 5,109,791<br>135,600<br>4,103<br>1,855 | 5,109,781<br>135,600<br>4,103<br>1,820 | 5,109,777<br>135,601<br>4,101<br>1,810 |

#### Supplementary Table 12: Assembly comparison using different filtering parameters: EC958 read and assembly accuracy (qcat demultiplexing)

| **Basecalling** | **Guppy3.4.3_hac** | | | | **Guppy3.6.1_hac** | | | |
|---|---|---|---|---|---|---|---|---|
| **Filtering parameter** | All reads (Porechop trimmed) | Japsa --lenMin 1000 --qualMin 10 | Japsa --lenMin 2000 --qualMin 5 | Filtlong --min_length 1000 --keep_percent 90 | All reads (Porechop trimmed) | Japsa --lenMin 1000 --qualMin 10 | Japsa --lenMin 2000 --qualMin 5 | Filtlong --min_length 1000 --keep_percent 90 |
| Number of reads | 239,123 | 202,759 | 175,104 | 138,826 | 242,794 | 205,223 | 176,916 | 139,921 |
| Average read percent identity | 91.0 | 90.8 | 90.8 | 91.0 | 93.7 | 93.6 | 93.5 | 93.7 |
| Mean read quality | 11.5 | 11.5 | 11.5 | 11.7 | 13.6 | 13.6 | 13.5 | 13.7 |
| Read length N50 | 13,106 | 13,249 | 13,557 | 14,393 | 13,042 | 13,181 | 13,489 | 14,321 |
| **Final assembly comparison** | | | | | | | | |
| Assembly nucleotide identity (%) | 99.99 | 99.99 | 99.99 | 99.99 | 99.99 | 99.99 | 99.99 | 99.99 |
| Number of SNP (DNAdiff) | 35 | 23 | 26 | 33 | 2 | 4 | 4 | 2 |
| Number of GSNP (DNAdiff) | 6 | 3 | 3 | 4 | 1 | 1 | 1 | 1 |
| Number of indels (DNAdiff) | 41 | 45 | 46 | 44 | 26 | 25 | 27 | 27 |
| Assembly quality score (Pomoxis) | 48.32 | 48.10 | 48.98 | 48.27 | 52.67 | 52.27 | 53.52 | 51.87 |
| Mismatches per 100 kb (QUAST) | 0.67 | 0.44 | 0.50 | 0.63 | 0.04 | 0.08 | 0.08 | 0.04 |
| Indels per 100 kb (QUAST) | 0.78 | 0.88 | 0.90 | 0.86 | 0.51 | 0.50 | 0.53 | 0.53 |
| Contig Size (bp) | 5,109,762<br>135,600<br>4,284<br>1,610 | 5,109,793<br>135,596<br>4,208<br>1,823 | 5,109,797<br>135,600<br>4,180 | 5,109,762<br>135,600<br>4,093 | 5,109,782<br>135,600<br>4,100<br>1,810 | 5,109,777<br>135,600<br>4,103<br>1,820 | 5,109,781<br>135,600<br>4,116 | 5,109,783<br>135,598<br>4,098 |

#### Supplementary Table 13: Assembly comparison using different filtering parameters: EC958 read and assembly accuracy (guppy demultiplexing)

| **Basecalling** | **Guppy3.4.3_hac** | | | | **Guppy3.6.1_hac** | | | |
|---|---|---|---|---|---|---|---|---|
| **Filtering parameter** | All reads (Porechop trimmed) | Japsa --lenMin 1000 --qualMin 10 | Japsa --lenMin 2000 --qualMin 5 | Filtlong --min_length 1000 --keep_percent 90 | All reads (Porechop trimmed) | Japsa --lenMin 1000 --qualMin 10 | Japsa --lenMin 2000 --qualMin 5 | Filtlong --min_length 1000 --keep_percent 90 |
| Number of reads | 227,542 | 192,927 | 166,613 | 132,076 | 238,139 | 201,130 | 173,392 | 137,395 |
| Average read percent identity | 91.1 | 91.0 | 91.0 | 91.2 | 93.8 | 93.7 | 93.6 | 93.8 |
| Mean read quality | 11.6 | 11.6 | 11.6 | 11.7 | 13.7 | 13.6 | 13.6 | 13.8 |
| Read length N50 | 13,102 | 13,240 | 13,547 | 14,387 | 13,031 | 13,173 | 13,480 | 14,301 |
| **Final assembly comparison** | | | | | | | | |
| Assembly nucleotide identity (%) | 99.99 | 99.99 | 99.99 | 99.99 | 99.99 | 99.99 | 99.99 | 99.99 |
| Number of SNP (DNAdiff) | 24 | 28 | 29 | 43 | 4 | 4 | 4 | 2 |
| Number of GSNP (DNAdiff) | 4 | 6 | 9 | 7 | 1 | 1 | 1 | 1 |
| Number of indels (DNAdiff) | 37 | 35 | 38 | 46 | 24 | 25 | 22 | 26 |
| Assembly quality score (Pomoxis) | 45.88 | 48.76 | 48.93 | 47.56 | 52.70 | 52.41 | 52.83 | 52.54 |
| Mismatches per 100 kb (QUAST) | 0.46 | 0.53 | 0.55 | 0.82 | 0.08 | 0.08 | 0.08 | 0.04 |
| Indels per 100 kb (QUAST) | 0.72 | 0.69 | 0.74 | 0.88 | 0.48 | 0.48 | 0.46 | 0.51 |
| Contig Size (bp) | 5,109,787<br>135,601<br>4,110<br>1,833 | 5,109,791<br>135,600<br>4,103<br>1,855 | 5,109,769<br>135,600<br>4,108 | 5,109,795<br>135,600<br>4,190 | 5,109,777<br>135,600<br>4,103<br>1,825 | 5,109,776<br>135,600<br>4,100<br>1,809 | 5,109,779<br>135,600<br>4,092 | 5,109,782<br>135,598<br>4,093 |

**Supplementary Figures:**

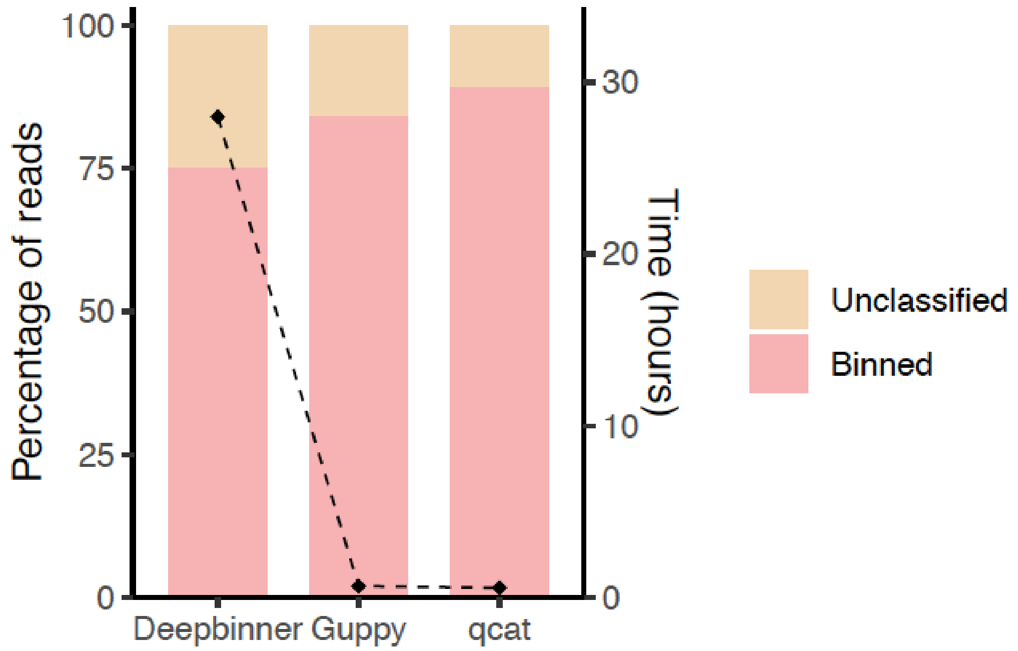


**Supplementary Figure 1: Demultiplexing metrics**

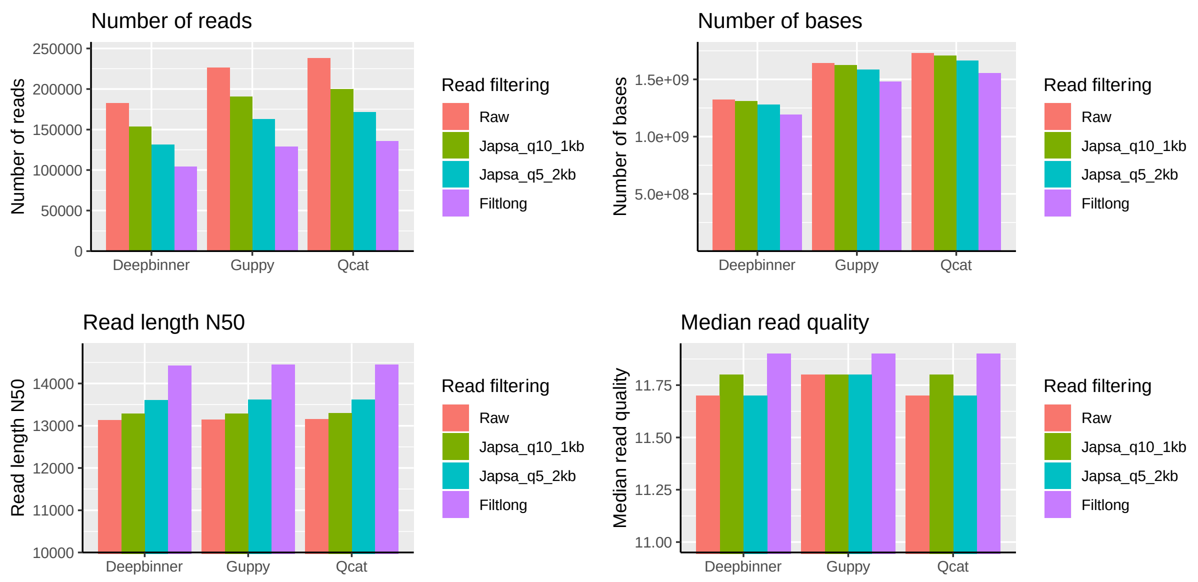


**Supplementary Figure 2: Long-read metrics using different demultiplexing tools and read filtering parameters (using EC958 ONT data)**

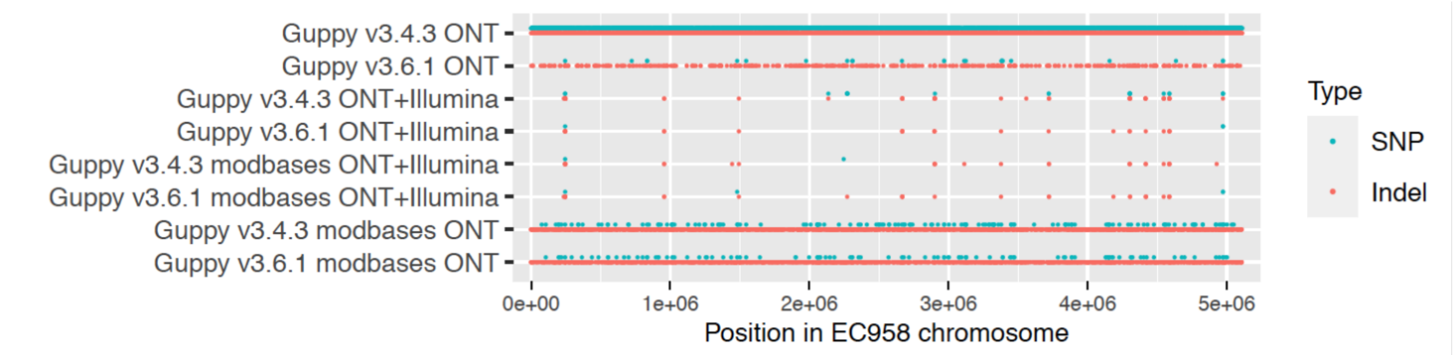


**Supplementary Figure 3: comparison of SNPs/indels in ONT assemblies vs. complete EC958 chromosome**

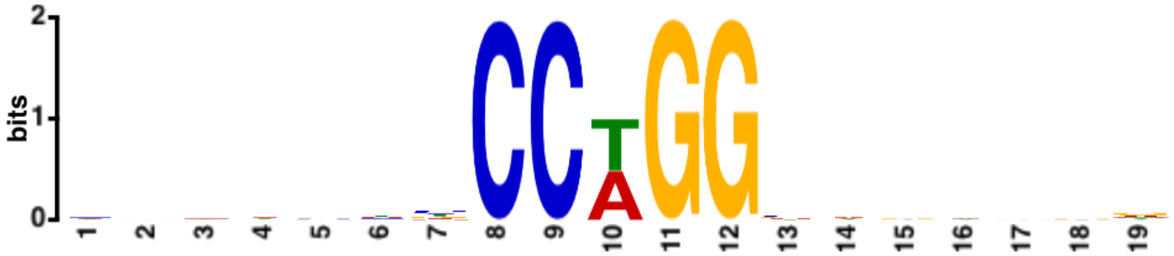


**Supplementary Figure 4: Motif enriched in the sequences around the 401 shared SNPs from the branch leading to discrepant ONT assemblies as indicated by the star in Figure 5A (main text).**
